# Transcriptional Pathology Evolves Over Time in Rat Hippocampus Following Lateral Fluid Percussion Traumatic Brain Injury

**DOI:** 10.1101/2021.04.29.442035

**Authors:** Rinaldo Catta-Preta, Iva Zdillar, Bradley Jenner, Emily T. Doisy, Kayleen Tercovich, Alex S. Nord, Gene G. Gurkoff

## Abstract

Traumatic brain injury (TBI) causes acute and lasting impacts on the brain, driving pathology along anatomical, cellular, and behavioral dimensions. Rodent models offer the opportunity to study TBI in a controlled setting, and enable analysis of the temporal progression that occurs from injury to recovery. We applied transcriptomic and epigenomic analysis, characterize gene expression and in ipsilateral hippocampus at 1 and 14 days following moderate lateral fluid percussion (LFP) injury. This approach enabled us to identify differential gene expression (DEG) modules with distinct expression trajectories across the two time points. The major DEG modules represented genes that were up- or downregulated acutely, but largely recovered by 14 days. As expected, DEG modules with acute upregulation were associated with cell death and astrocytosis. Interestingly, acutely downregulated DEGs related to neurotransmission mostly recovered by two weeks. Upregulated DEG modules related to inflammation were not necessarily elevated acutely, but were strongly upregulated after two weeks. We identified a smaller DEG module with delayed downregulation at 14 days including genes related to cholesterol metabolism and amyloid beta clearance. Finally, differential expression was paralleled by changes in H3K4me3 at the promoters of differentially expressed genes at one day following TBI. Following TBI, changes in cell viability, function and ultimately behavior are dynamic processes. Our results show how transcriptomics in the preclinical setting has the potential to identify biomarkers for injury severity and/or recovery, to identify potential therapeutic targets, and, in the future, to evaluate efficacy of an intervention beyond measures of cell death or spatial learning.

## INTRODUCTION

The U.S. Center for Disease Control and Prevention (CDC) reported almost 3 million traumatic brain injury (TBI) incident-related in- and outpatient emergency room visits in the U.S. in 2014 ^1^. Among military service personnel, almost 20,000 soldiers experienced a TBI in 2019, mostly mild to moderate ^2^. While severe TBI can have long-lasting effects resulting in a chronic disease state ^3^, most individuals who experience a mild, and many with a moderate severity of TBI do recover. However, it has become clear that even mild and moderate TBI are associated with increased risk for late-onset neurodegenerative diseases ^4–7^ such as Parkinson’s and Alzheimer’s disease. Resolving the longitudinal molecular and cellular changes associated with TBI is critical towards understanding the etiology driving acute and lasting health impacts.

Measurement of the perturbations in gene expression in bulk tissue and single cells following TBI in model systems has provided a systems-level perspective of the impacts of TBI ^8–12^. In the initial period following TBI, transcriptional studies have identified signatures of neuroinflammation, and cell death, that correlate with long-term outcomes such as cognitive dysfunction. In contrast to the better-characterized initial response, longer-term transcriptional impacts have not been as well studied. Parallel to transcriptomic changes, initial studies of the epigenetic impacts of TBI ^8,13,14^ found that 7 days post-injury, DNA methylation was impacted at hundreds of sites across the genome. Other epigenetic signatures, for example changes to histone proteins, have also been reported after TBI ^13,15^. Taken together, previous transcriptomic studies have contributed to a systems-level understanding of acute and long-term TBI-associated molecular pathology. However, these single time-point studies do not directly elucidate the progression of changes linking injury to recovery or highlight stage-specific biomarkers that can serve , e.g. as markers for recovery state.

Changes in gene expression following TBI are dynamic and responsive to the state of the recovery following injury. For example, it is well described that cell death peaks acutely post-injury, and that there are immediate changes in neurotransmission that reduce plasticity ^16,17^. Thus, a time course transcriptomics approach has the potential to resolve the temporal progression of TBI pathology, as well as to discover new candidate markers and pathways. Finally, linking transcriptional and epigenetic changes can illuminate genomic mechanisms underlying long-term molecular pathology. Towards these goals, we interrogated acute (1-day) and subchronic (14-day) changes in ipsilateral hippocampus in rats exposed to Lateral Fluid Percussion (LFP) injury. Our results map molecular signatures associated with distinct expression trajectories following TBI and find evidence of parallel alterations in histone H3K4me3 at relevant gene promoters.

## RESULTS

### Gene Expression Perturbations Following TBI are Both Transient and Persistent

Adult male Sprague-Dawley rats were randomly assigned to sham control (n = 6) and LFP (n = 8) groups (Figure 1A). These groups were separated for acute assessment one day post-injury (n = 2 sham, n = 4 LFP) and subchronic assessment on day 14 post-injury (n = 4 sham, n = 4 LFP). RNA sequencing (RNA-seq) was performed on bulk ipsilateral hippocampal tissue. We determined differential gene expression using multiple-testing corrected p-value < 0.05 between TBI and sham control at each time point. Full differential gene expression results are reported in Tables S1-S3. Overall, we identified 1351 up- and 1013 downregulated DEG 1 day following TBI, and 439 up- and 41 downregulated DEGs at 14 days, relative to timepoint-matched sham controls. Sham and TBI samples hierarchically clustered by group and timepoint, indicating overall DEG signatures that robustly discriminate TBI from sham as well as acute as compared to subchronic time points (Figure 1B). A volcano plot representation of expression changes and the magnitude DEG significance for all tested genes, with top DEGs (lowest p-value and/or highest absolute expression fold change) labeled (Figure 1C). Finally, Principal Component Analysis (PCA) using full transcriptomic data similarly captured well-defined separation of condition and time point (Figure 1D), with the 1-day TBI signature driving the largest proportion of variance in PCA space (i.e. PC1).

**Figure 1.**
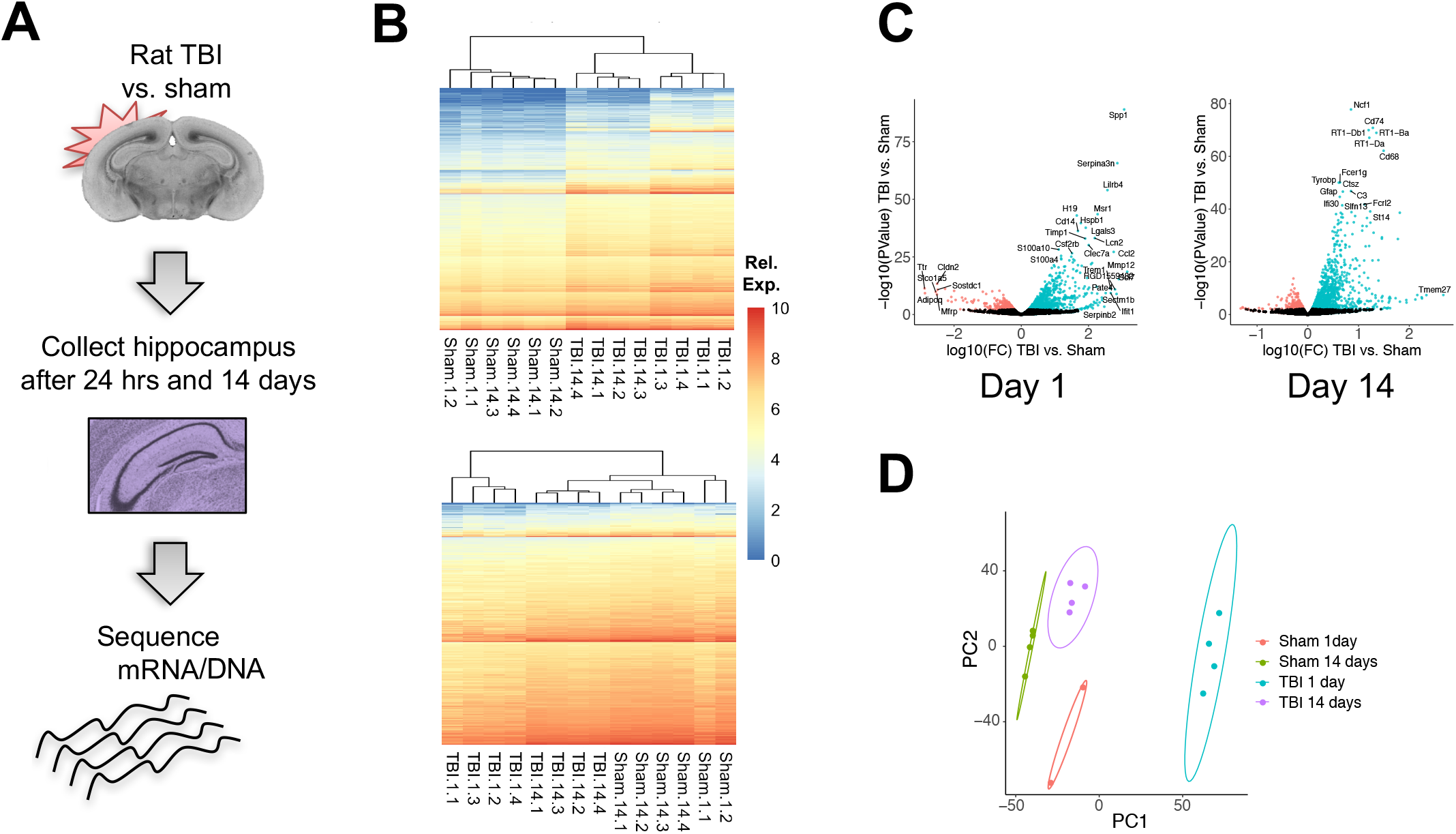
Differential Expression of Genes Subjected to TBI. (A) Summary of experimental design. (B) Clustered heatmap of reads from up- (upper panel, n = 6,870) and downregulated genes (lower panel, n = 10,302) at both time points (1 and 14 days after TBI) for genes with un-adjusted p-value < 0.05. (C) Volcano plots of the differentially expressed genes at each of the time points. Each dot represent a gene; those colored in red were downregulated, while the ones in green were upregulated, TBI vs. sham control (significance level is 99%, *α* = 0.05). Differential gene expression presented in log10 scale, with example genes labeled. (D) Principal Component Analysis (PCA) plot showing the first two components. Ellipses represent cluster confidence intervals at the 95% confidence level (when there were only two data points, an ellipse was manually drawn to enclose both points).

Genes that were differentially expressed at adjusted p-value < 0.05 at either time point (4354 total genes) were assigned to a co-expression module based on expression changes across time points (Figure 2A). A relatively small subset of genes were increased early after injury and then further increased in expression at 14 days (80 genes, module i.). A larger subset had a larger increase on day 1 as compared to 14, but remained significantly elevated at the later time point (238 genes, module ii.). The largest module of genes (n = 2085) were significantly upregulated at 1 day but had recovered by 14 days as compared to shams from the 14-day time point (module iv.). This was followed in size by the module of genes that showed the opposite effect of downregulated at day 1 but recovered by 14 days (1789 genes, module v.). Only 22 acutely downregulated genes demonstrarted only partial recovery (module vii.) Finally, 9 acutely downregulated genes were even more significantly perturbed at 14 days (module viii.). In addition to genes that were altered on the first day post-injury, there were two additional subsets of genes that were not acutely changed, but had differential expression on day 14. Of these genes, 113 had delayed upregulation (module iii.) and 18 were downregulated only at day 14 (module vi.). Overall, this analysis identified clear differences in expression and recovery among genes sensitive to TBI, which may reflect both the trajectory of recovery and the onset of long-term pathology.

**Figure 2.**
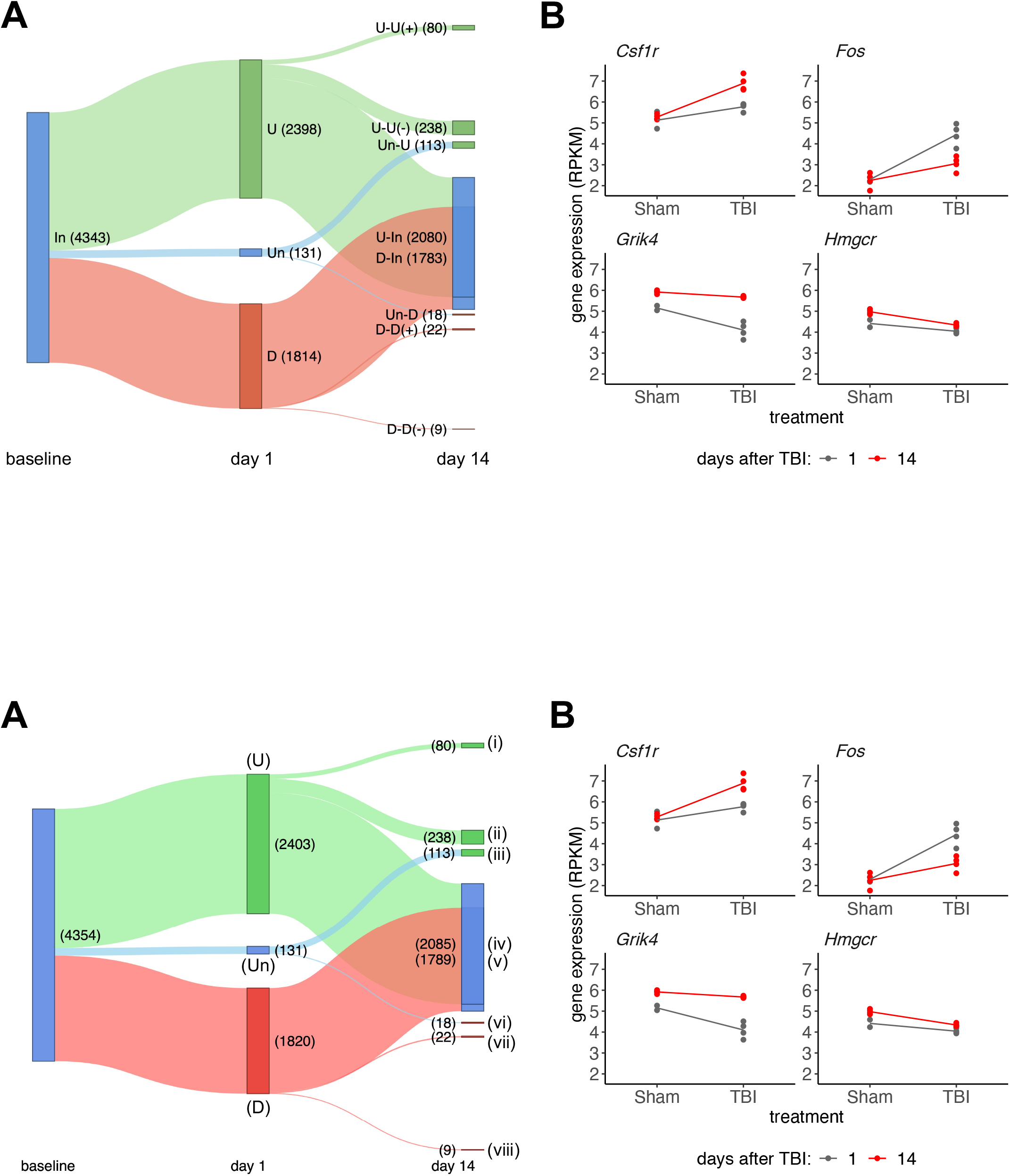
Gene Expression Trajectories Over Time After TBI. (A) Sankey plot showing the relative gene expression trajectory across the period 1 to 14 days after TBI (limma-voom model with p <= 0.05, expression normalized/corrected for batch effect and sham effect over time). Gene trajectory modules are labeled as follows: (i) denotes persistently upregulated genes; (ii) means acutely upregulated genes showing partial recovery at day 14; (iii) denotes genes with delayed upregulation; (iv) and (v) denote acutely up- and downregulated genes, respectively, showing full recovery at day 14; (vi) represent genes with delayed downregulation; (vii) depicts acutely downregulated genes showing partial recovery; and (viii) denotes persistently downregulated genes. (B) Expression level comparisons at both time points, TBI and sham control, for selected genes (*Csf1r*, *Fos*, *Grik4*, and *Hmgcr*) showing different trajectories. Gene expression is given in log RPKM (Reads Per Kilobase of transcript, per Million mapped reads).

### DEG Expression Trajectory Modules are Associated with Distinct TBI Pathology

Many DEG are related to known changes following injury, the trajectory module for such genes can give insights into temporal and biological processes associated with TBI. For example, Figure 2B shows expression differences for four example genes with known TBI associations: *Csf1r* (neuroinflammation, module iii), *Fos* (persistent activation, module iv), *Grik4* (synaptic signaling, module v), and *Hmgcr* (cholesterol synthesis, module vii). Moving from known markers to a systems level approach to understand trajectory modules, we next sought to understand which biological pathways and processes are associated with each of the differential expression trajectories via Gene Ontology (GO) analysis performed on module DEG sets (p-value and q-value cutoffs of 0.05; full results in Table S4, Figure 3A). As expected, the modules harboring DEGs noted above were associated with expected biological functions. In addition to these representative genes and pathways, we report the set of enriched GO terms for each trajectory module and show representative DEGs in Figures 3B-G.

**Figure 3.**
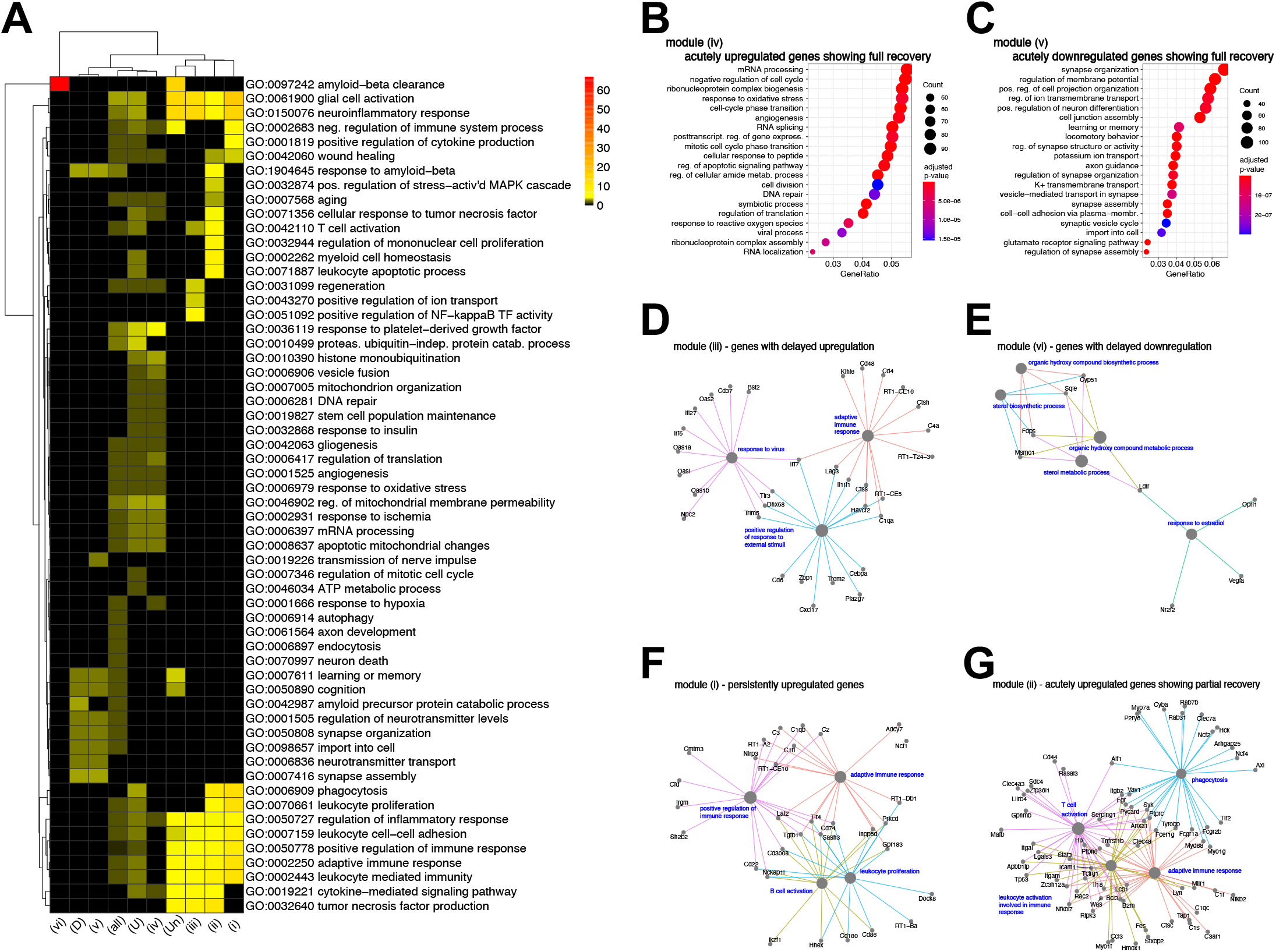
Gene Ontology (GO) Analysis Stratified According to Expression Trajectories. (A) Heatmap depicting brain-specific or general biological term enrichment for clusters identified as in Figure 2A (adjusted p-value <= 0.05, 10 top enriched terms for each gene group defined). (B) Dot plot showing significance of top 20 GO categories for modules showing acutely upregulated genes showing full recovery at day 14 (p-value <= 0.05, q-value <= 0.05). (C) Same as (B) for acutely downregulated genes. (D) Network plot depicting genes annotated to up to the 5 most significant GO terms for the module with delayed upregulation of gene expression in response to TBI (only brain-related or related terms shown). (E) Same as (B) for the module with delayed downregulation of gene expression. (F) Same as (B) for the module showing persistent, increasing perturbation in gene expression. (E) Same as (B) for the module showing acutely upregulated genes showing partial recovery at day 14.

Most of the GO terms that were enriched acutely (Figure 3B, module iv) were associated with biological processes similar to previously described physiological progression of TBI recovery ^18^. As examples, *Capn2* and *Bax*, both involved regulation of apoptotic cell death, were expressed acutely, but returned to baseline by day 14. Similarly, well-characterized inflammatory markers such as *Il1b* and *Tnf* were induced acutely and then returned to baseline by day 14. Other module iv upregulated GO terms included cellular homeostasis and immune/angiogenic response (RNP complex biosynthesis, angiogenesis, DNA repair, cell division, etc.), immediate response to environmental stimulus (response to oxidative stress, cellular response to peptide, etc.), and apoptosis (Figure 3B, module iv.). Module v, representing genes that were acutely downregulated, was enriched for GO terms associated with synapse function and organization, potentially capturing changes in neuronal state and communication in response to the acute excitotoxic period and the activation of surrounding astro- and microglia (Figure 3C, module v.). Over half of the DEGs identified as acutely altered following TBI were included in these two modules and, as described above, the majority return to pre-injury levels by 14 days. Genes in these acute modules (e.g., *Ddc*, *Cdhr1*) overlap with transcriptomic studies reported 7 days after TBI in a similar animal model ^8^.

Of genes that are differentially expressed at 14 days, many were also differentially expressed acutely. These include genes in the module ii that were upregulated at both time points, which were enriched for GO terms associated with immediate and adaptive immune response, and phagocytosis (Figure 3G). As a representative gene, *Il18* is an cytokine of the IL-1 family involved in neuroinflammation ^19^. Another example, *Tyrobp* has previously been identified as chronically upregulated following TBI ^20^. In contrast to the relatively large number of acutely upregulated genes showing partial recovery, only 22 acutely downregulated genes (module vii) showed continued downregulation at day 14, and these genes did not show statistically significant enrichment of specific GO terms.

While there were a relatively small number of DEGs (80) in module i, where expression continued to increase at day 14, these genes were strongly associated with specific GO terms, such as glial cell activation. Likewise, very few genes (9) exhibited increased downregulation at day 14, with no enrichment of brain-specific GO terms. The final modules represent genes that had a pattern of delayed transcriptional perturbation, with expression similar to 14-day sham on day 1 post-TBI but significantly different after two weeks. GO terms associated with delayed upregulation (module iii, 113 DEGs) are associated with innate and adaptive immune activation and inflammation (Figure 3D). As an example, *Nrlp3* has been shown to be a driver of inflammation and immune response following TBI ^21^. A very limited number of DEG (18) exhibited a delayed downregulation (module vi), and were associated with sterol biosynthesis and metabolism (Figure 3E), suggesting a possible change in cell membrane-related cholesterol homeostasis ^22^.

### Parallel H3K4me3 Epigenetic Changes at DEG Promoters One Day After TBI

To explore the link between transcriptional changes and TBI-induced acute epigenetic effects, we conducted chromatin immunoprecipitation with sequencing (ChIP-seq) with antibodies targeting H3K4me3, a histone H3 post-translational modification found to be a signature for active transcription history that is characteristically found at gene promoters ^23,24^. We compared ChIP-seq coverage in TBI samples versus sham controls at each gene promoter, and across promoters associated with DEGs and their expression trajectory modules. H3K4me3 is generally associated with transcriptional activation, and, as expected, overall gene expression levels were correlated with H3K4me3 at the promoter.

In line with DEG changes following TBI, upregulated genes had generally higher H3K4me3 enrichment in TBI versus sham samples and the inverse was true for down-regulated genes (Figure 4A). Changes in H3K4me3 in the TBI samples were generally subtle, and many DEGs did not exhibit changes at the level of sensitivity of these experiments. However, a small set of upregulated DEGs (n = 223) showed strong enrichment of H3K4me3 in TBI samples when compared with sham controls, as indicated in the dashed box in Figure 4A. For these genes, the level of promoter H3K4me3 in sham controls was low or undetectable, suggesting transition from silent to active transcriptional state following TBI. The genes exhibiting this pattern were highly enriched in GO terms associated with leucocyte migration and activation of immune response and phagocytosis (Table S5). While most DEG exhibited subtle epigenetic impacts, we found significant increases in H3K4me3 for all but one cluster of acutely upregulated expression trajectory modules (p < 0.05) and a significant decreases H3K4m3 for acutely downregulated modules (Figure 4B). Figures 4C and 4D depict ChIP-seq coverage genomic representation of H3K4me3 changes for TBI and sham controls at day 1 around up- and downregulated DEGs, respectively. Genes shown are *Bcl3*, *Scocs3*, and *Fos* for upregulated, and *Car2*, *Tgfb2*, *Ank3* for downregulated.

**Figure 4.**
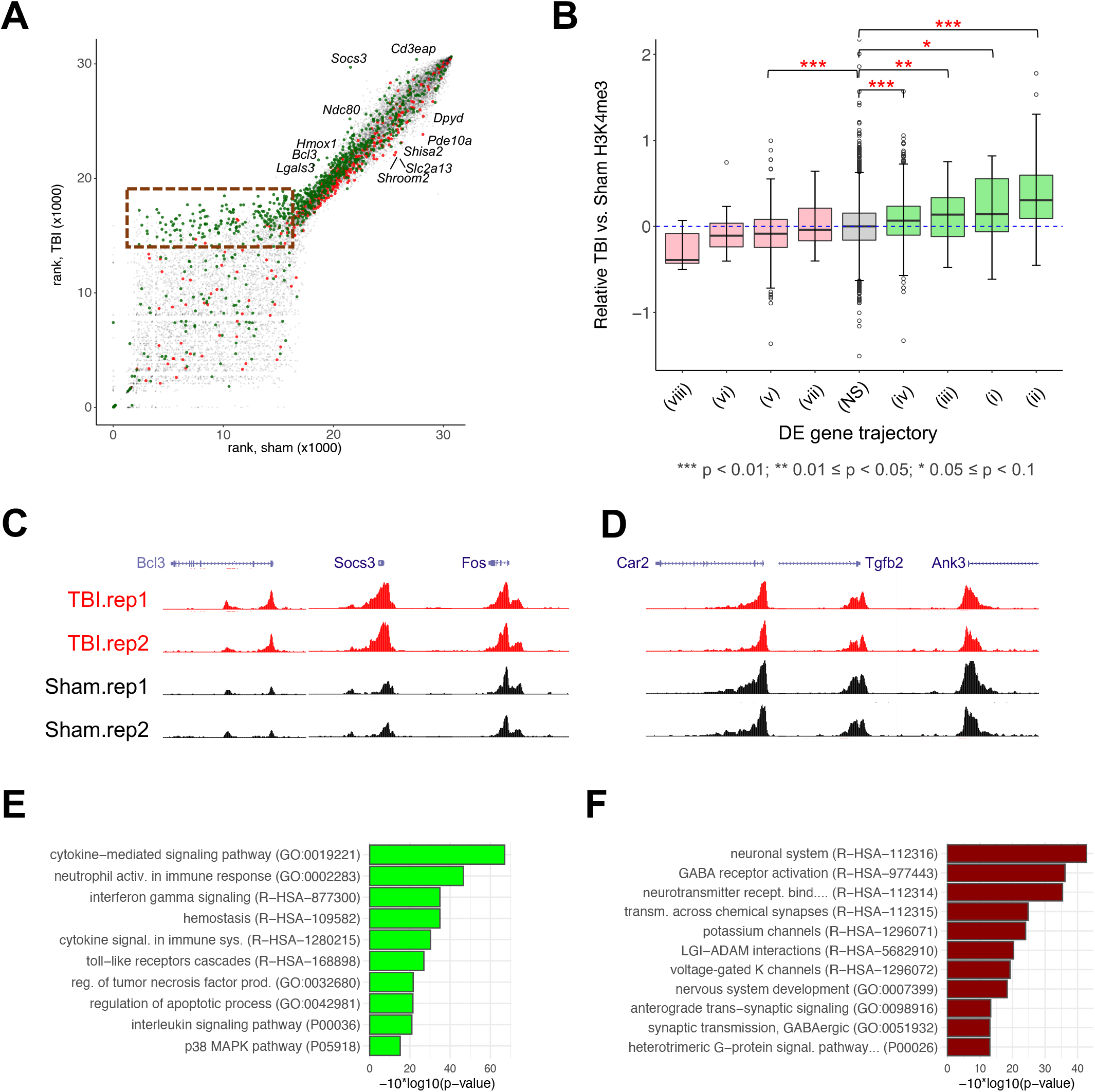
H3K4me3 at Promoters of Genes Differentially Expressed after TBI. (A) Coverage rank plot for TBI versus sham for H3K4me3 at day 1. Y-axis is for the TBI sample, while the x-axis is for the sham control. Each dot is a gene promoter locus present at either of the two conditions. Green-colored dots are associated with promoters of upregulated genes, while red-colored ones are for downregulated genes. Region highlighted with dashed red box denotes genes with distinctive H3K4me3 enrichment in TBI at day 1. Dots labeled with name of genes were manually annotated for example genes that showed more prominent differences between TBI and its sham control. (B) Box plot showing the distribution of relative likelihood ratios for each of the DE gene trajectories defined in Figure 2A compared to genes that were not differentially expressed (NS). Green-and pink-filled boxes denote up- and downregulated associated genes, respectively; Red stars above the plot mean the statistical significance of Tukey means comparison between groups and the NS group (p-values shown below the plot). (C) Example of a representation of H3K4me3 genomic enrichment coverage region for sham and TBI at day 1 for upregulated loci (*Bcl3*, *Socs3,* and *Fos*). (D) Same as (C) for downregulated genes (*Car2*, *Tgfb2*, and *Ank3*). (E) Bar plot depicting the most significant function annotation terms (GO biological process, Reactome and PANTHER pathways) for the upregulated genes showing H3K4me3 differential enrichment. (F) Same as (E) for downregulated genes.

Gene Set Enrichment Analysis (GSEA) was conducted on the set of 100 up- and downregulated genes with strongest acute change in H3K4me3. As noted above, genes with largest increases in H3K4me3 were associated with inflammation/immune response and are acutely upregulated genes showing partial recovery at day 14. Overall, genes with strong epigenetic changes at day 1 were enriched with GO terms and biological pathways akin to the full set of up- and downregulated DEGs, reflecting the same general pathophysiological processes are associated with transcriptional and epigenetic response to TBI. These findings indicate that epigenetic changes in H3K4me3 both parallel and can precede transcriptional changes, as all trajectory modules showed some evidence for epigenetic changes in the same direction as transcriptional changes.

## DISCUSSION

TBI activates a cascade of events that can lead to acute secondary effects as well as chronic pathology. In the LFP model, cell death peaks in the first 24 hours, with negative effects significantly reduced by one week following injury ^25^, and cognitive performance is most significantly diminished in the first weeks following injury ^26^. Similarly, well-characterized is the acute inflammatory response and attenuated neuroinflammation often persists for the duration life ^27^ and is linked to neurodegeneration ^28,29^. Towards understanding the progression of pathology following TBI, we evaluated changes in hippocampal gene expression acutely (1 day) following injury and after 14 days. Our results not only corroborate previous acute transcriptome studies, but also recapitulate known physiological pathology that occurs in the injury and recovery ^30,31^. Our findings reflect the excitotoxicity peak acutely post-injury, with DEGs and GO terms associated with ongoing cell death mostly absent delayed time point. Somewhat surprisingly, while downregulation of neurotransmission and synaptic plasticity genes were also obvious acutely, these genes were included in modules that largely resolved by 14 days. Our results capture neuroinflammation responses that separate into DEG modules that peak acutely, that maintain similar elevation in the transition from acute to two weeks out, and that are higher at 14 days post-injury. Finally, we identified a delayed onset DEG module comprised of genes related to cholesterol metabolism and amyloid-beta. This last signature is of particular novelty and interest, as formation of plaques in the rat model of LFP has not been observed. Finally, we identified epigenetic substrates of DEG evident at 1 day post-injury. TBI-induced genes exhibited promoter H3K4me3 changes, indicating epigenetic activation as a response to TBI and suggesting that such changes may underlie long-term transcriptional dysregulation, particularly of inflammation and neuroimmune activation genes.

There are many well described immediate physiological impacts of injury including the permeabilization of the blood-brain barrier (BBB), axonal shearing, and toxic neuronal hyperexcitability ^18,32,33^ that occur acutely following injury. Changes in pathways related to apoptosis, necrosis and synaptic plasticity are also captured in the first hours following TBI ^30,31^. In response to primary injury, leukocytes, macrophages, and lymphocytes enter from the blood stream and secrete various cytokines enhancing the inflammatory response in the injured site ^34^. In addition, the increased porosity of the BBB after TBI permits extensive extravasation of plasma proteins onto the brain, which further induces inflammatory responses due to the neurotoxicity of those molecules ^35^. In response to this acute neurotoxicity microglia are activated. The sum of these changes contributes to edema and can lead to the eventual formation of a cystic cavity and activation of glial cells around the lesioned tissue. Overall, these well-characterized acute pathophysiological responses to TBI were reflected in significant changes in DEG signatures and related GO terms observed 24 hours following moderate fluid percussion injury.

While other systems-level analysis have similarly identified transcriptomic signatures associated with acute processes, having the ability to compare changes at one and 14 days following TBI resolves how immediate post-injury responses progress over time. Enrichment for apoptosis terms was highly significant in acutely upregulated genes showing full recovery at day 14. DEG (module iv), supporting findings that by 72 hours post-injury, while cell death continues, it is significantly diminished ^25^. Similarly, genes associated with regulation of synaptic plasticity (GO:0048167) were represented in the module of acutely downregulated genes showing full recovery. Genes in this category included *Adcy8*, *Bcan*, *Camk2b*, *Neurod2*, *Slc8a2*, and others (Table S4), and are part of other terms as well, e.g. learning or memory (GO:0007611). One of our hypotheses was that genes related to synapses, neurotransmission and plasticity would be maximally decreased acutely following injury, but would remain significantly downregulated out to two weeks. Thus it was surprising that while we detected systems-level changes in synaptic DEGs and GO terms acutely, values returned to baseline by day 14. Thus, our results suggest that the large scale transcriptional perturbation is not persistent. However, it is possible that transcriptional perturbations remain altered in a subset of cells, and are not captured given the volume and diversity of the tissue evaluated. It is also possible that cell death has long term impacts to synaptic function and plasticity that can’t be rescued even if transcription is more appropriately regulated in the surviving neurons. Further, changes in protein levels do not always follow changes in mRNA expression; in particular during periods of dynamic transitions ^36^. Therefore, even if transcription returns toward baseline, there could be persistent protein dysregulation or changes in synaptic structure and function triggered acutely that explain the lasting deficits in neurotransmission, long-term potentiation and behavior.

Our data demonstrate the power of time course evaluations to chart the course of transcriptomic sequalae caused by experimental TBI. Our findings recapitulate several previously published observations ^5,8,11,12,37^. Of note, the signatures identified here differ substantially between the genes that are induced (i.e. upregulated) following injury and those that are depressed. Moreover, TBI appears to induce waves of primarily (modules ii and iv) and secondarily (module iii) induced genes corresponding to stages of inflammation and immune response that are observed both acutely 1 day following injury as well as two weeks later. In contrast, downregulation was dominated by early changes at day 1 with most genes returning to baseline by day 14 and only a small, but relevant, set of new DEGs emerging at this later time period. Many of the chronically downregulated DEGs are involved in processes relevant to recovery and long-term neurodegeneration following TBI. As TBI can result in progressive inflammation and neurodegeneration and lasting behavioral disorders, intersecting transcriptomics and epigenetic changes as the brain transitions from an acute response to recovery and ultimately into the chronic state can provide insight into long-term genomic impacts. Our, H3K4me3 ChIP-seq data indicates that TBI-induces parallel transcriptomic and epigenetic changes overall, and that genes involved in neuroinflammation show the strongest chromatin response at 24 hours, potentially priming these pathways for long-term epigenetic activation.

Some of the more interesting findings related to genes that had a delayed response to injury. For example, amyloid-beta clearance was a highly enriched GO term in the delayed downregulation group (Figure 3A). In addition, a small set of loci associated primarily with sterol metabolism and maintenance were significantly decreased only at the delayed 14-day time point. TBI and cholesterol metabolism have been suggested as causative factors in Alzheimer’s and other neurogenerative diseases ^38–41^. There was also a small number of genes (n = 9) that were downregulated early, but were even further diminished at the two-week time point including *Adra2a*, *Ccdc33*, *Dpp10*, *Fdft1*, *Hmcn1*, *Klhl14*, *Slc38a4*, *Tacr3*, and *Tfrc*. Each of these genes have been reported as downregulated in models of chronic neurodegenerative disease such as Alzheimer’s and Parkinson’s ^41,42,51,52,43–50^.

This study presents a systems-level and genomic perspective to understanding acute and long-lasting effects of TBI. Critically, doing a longitudinal study including a two week outcome identified altered recovery of up- and dow-regulated genes, differentiated acture and sub-chronic changes, separated immune and inflammation among those that resolve versus persist, and identified delayed upregulation in genes related to neurodegenerative disorders. Our findings also link epigenetic activation to acute and lasting transcriptional changes, highlighting strong epigenetic activation of immune and infkammation genes. These data demonstrate the power of longitudinal epigenetic and transcriptomic analyses to understand mechanistic changes in the days-to-weeks following TBI. Future studies are needed to model the time-course at finer scale, including longer-term changes in expression and genome state. These studies should necessarily evaluate not only change in transcription, but also protein expression. Moreover, our focus on ipsilateral hippocampus should capture direct impacts of injury, but there could be systematic impacts across the brain, and future analyses should consider other regions of the brain that may be directly affected by injury as well as distal areas that are presumed to be healthier. Finally, expanded single cell studies paired with spatial analysis have the potential to resolve systems-level pathology identified in bulk studies such as ours, to specific cells and circuits.

## CONCLUSION

Ultimately, there are two major opportunities for the future use of findings here and in other similar studies. First, to identify genes that can be used to generate prognostic and theragnostic tools for physicians, to determine stage and severity of injury and recovery and to distinguish which individuals are more likely to develop long-term dysfunction and when a drug might be exerting expected beneficial effects. Second, time course transcriptomic datasets such as ours will help to identify potential targets for therapeutic intervention, not only in the acute hours-to-days post-injury during the in-hospital setting, but also remote the initial recovery period after the patient has returned home. In addition to revealing underlying neurobiology of the progression of TBI-associated transcriptional pathology, the TBI-associated genes annotated to trajectory modules here represent a valuable resource for the field regarding time-dependent biomarkers and candidates for in-depth study.

## ACKNOWLEDGEMENTS

ASN and RC-P were supported by UC Davis institutional support to ASN and NIH NIGMS R35GM119831. RC-P was supported by a Science without Borders Fellowship from CNPq (Brazil). ETD, KT and GG were supported by UC Davis institutional support and NINDS R01NS084026.

## AUTHOR CONTRIBUTIONS

Conceptualization: RCP, ASN, and GG; Methodology and Investigation: ETD (surgery), KT (sample collection), IZ (RNA-seq, ChIP-seq), RCP (bioinformatics, modelling), BJ (bioinformatics); Software: RCP and BJ; Writing – Original Draft: RCP, ASN, and GG; Writing – Review & Editing: RCP, ASN, and GG; Funding Acquisition: GG and ASN; Supervision: ASN and GG.

## MATERIAL AND METHODS

### Test Animals

Adult male Sprague-Dawley rats (300-375g; Envigo, Livermore, CA, USA) were randomly assigned to sham control (n = 6) and LFP injury (n = 8) groups (Figure 1A). These groups were further randomly separated for acute assessment one day post-injury (n = 2 sham, n = 4 LFP) and chronic assessment on day 14 post-injury (n = 2 sham, n = 2 LFP). Rats were housed in a campus vivarium with regulated temperature (22 °C) and humidity (40-60%) and on a 12 hr light/dark cycle. Rats had free access to food and water throughout the experiment. All procedures adhere to the National Institutes of Health guidelines and were approved by the University of California, Davis Institutional Animal Care and Use Committee.

### Lateral Fluid Percussion (LFP) Injury Rat Model

Rats were randomly assigned to receive either LFP or sham injury ^53^. Sham animals received identical surgical procedures as TBI, including duration of anesthesia, except the fluid percussion injury was not administered. Anesthesia was induced using 4% isoflurane (in air). Animals were then intubated, shaved, transferred to a stereotaxic frame, and mechanically ventilated to maintain a surgical plane of anesthesia using 1.5 - 3% isoflurane (in 2 NO_2_ : 1 O_2_) for the remainder of the surgery. After sterile preparation, a subcutaneous injection of 0.25% bupivacaine (0.1 mL) was delivered to the shaved skin on the dorsal surface of the skull. A midline scalp incision was made, and the skin was retracted to expose the dorsal cranial surface. A circular 4.8 mm diameter parasagittal craniectomy was made over the right hemisphere midway between bregma and lambda and 3 mm lateral to midline using a trephine. Two stainless steel screws (0-80”) were secured in the skull, anterior and posterior to the craniectomy. A custom designed plastic tube injury hub was placed in the craniectomy and cemented to the skull with a combination of super glue gel and dental acrylic. The injury hub was then filled with sterile saline.

The fluid percussion device was calibrated to produce an injury of ~2.1 atm of pressure. In our hands, this injury typically results in a moderate injury with persistent spatial learning deficits for at least two weeks post-injury ^54,55^. The animal was then removed from anesthesia, attached to the device, and the injury hammer was released ^56^. Immediately following injury, animals were observed for the return of the toe pinch withdrawal, and then were returned to 2.5% isoflurane. Finally, the injury hub was removed, and the wound was sutured closed (4.0 braided silk suture), and the animal was placed in a heated cage and observed until they became sterile.

### Hippocampal Tissue Collection

Rats were placed in a chamber with continuous flow of 4% isoflurane with air as the carrier for 4 minutes. They were then removed from the chamber and rapidly decapitated. The brain was exposed and rolled out of the skull and placed on filter paper atop a glass dish sitting on ice. The cortex was hemisected rostral-to-caudal, then and rolled back (medial to lateral) exposing the underlying hippocampus. A curved, plastic spatula was used to roll the hippocampus out of the cortex and onto the filter paper. The same spatula was used to completely separate the hippocampus from the cortex and place it into a sterile microcentrifuge tube. Right and left hippocampus were collected separately.

### RNA-seq Experiments

Fresh hippocampus samples from the injury ipsilateral tissue were prepared as previously described ^57^. Total RNA was isolated from the samples using Ambion RNAqueous (Thermo Scientific cat # AM1912) and assayed using an Agilent BioAnalyzer instrument. We used TruSeq Stranded mRNA kits (Illumina P/N 20020594) to prepare the stranded mRNA libraries, that were sequenced on the Illumina HiSeq platform using a single-end 50-bp strategy. Libraries were pooled at 6-12 samples per lane; each library was quantified and pooled before submission for sequencing. All samples were sequenced at the UC Davis DNA Technologies Core.

### ChIP-seq Experiments

ChIP-seq was performed following established protocols ^58^. Frozen hippocampus tissue samples were individually crosslinked with a 1% formaldehyde buffer solution for 10 min, then washed and isolated. The resuspended crosslinked pellets were treated with protease inhibitor and sheared by sonication. Samples were then washed off of unbound DNA and free proteins, and incubated with Histone H3K4me3 antibody (mAb, Active Motif cat # 61979) and magnetic beads (Dynabeads) for 2 hr at 4 °C. After reaction, beads were magnetically separated, washed and de-crosslinked. The resulting DNA was then purified, size selected, and libraries were. prepared using Ovation Ultralow System V2 preparation kit (NuGEN P/N 0344). Input control libraries were prepared from DNA before antibody pulldown. The libraries were quantified and pooled prior to submission for sequencing. The UC Davis DNA Technologies Core sequenced the libraries as 50-bp single-end reads on an Illumina HiSeq 4000 instrument.

### Quantification and Statistical Analysis

#### Differential Gene Expression Analysis

We determined the differential gene expression between the TBI and sham control samples by individually aligning the FASTQ reads from the RNA-seq experiment to the rat genome (rn5) using STAR version 2.4.2a ^59^, after quality control evaluation using FASTQC version 0.11.8 ^60^ and counting reads with featureCounts version 1.5.0 ^61^, with UCSC gene annotations. We then analyzed the samples using a custom R script running the limma-voom model ^62^. For gene expression trajectories across 1 and 14 days, we established sham at day 14 as the baseline, and normalized TBI gene expression (RPKM) using the differential gene expression (DGX) determined by the limma-voom method. DEGs with similar trajectories were assigned into clusters, as depicted in Figure 2A. Baseline state was followed by the 1-day clusters: Upregulated (“U”), Unchanged (“Un”, not differentially expressed), and Downregulated (“D”). At 14-day, clusters were (i), (ii), and (iv) for upregulated genes at 1-day that were further upregulated, downregulated but not returning to baseline, and downregulated returning to baseline at day 14, respectively. Moreover, clusters labeled as (viii), (vii), and (v) were for downregulated genes at 1-day that were further downregulated, upregulated but not returning to baseline, and upregulated returning to baseline at day 14, respectively. Finally, (iii) and (vi) refer to cluster of genes that were not differentially expressed at 1-day (p < 0.05) but were up- and downregulated at 14-day, respectively. We conducted gene ontology analysis using the R clusterProfiler package ^63^ using p-value and q-value cutoffs of 0.05, for the genes in each of the expression pattern clusters defined at day 1 and day 14.

#### ChIP-seq Data Analysis

We quality-checked and trimmed adapter sequences from the reads of the FASTQ files from ChIP and input control samples using the FASTQC and Trim Galore! Version 0.5.0 ^64^, respectively. The filtered reads were aligned to the rn5 genome (UCSC gene annotation) with bwa version 0.7.16a ^65^, with duplicate reads removed with Samtools version 1.8 ^66^. We generated read coverage genome wide estimates and peak calls for sham and TBI samples using the pileup outputs from MACS2 version 2.1.2 ^67^. Using custom R scripts, we intersected coverages (pileups) from H3K4me3 in TBI samples and respective sham controls, and filter those annotated to gene promoters. For each individual set (TBI and sham), we ranked the coverages and used the ranks to compare against one another (to compensate for variations in sensitivity among data sets). We assigned up- or downregulated gene associated each TBI-sham pair at each time point by association with the differential expression data. We used MACS2 bdgdiff function to compare normalized H3K4me3 enrichment, focusing analysis on gene promoters. Log likelihood of differential H3K4me3 between TBI and sham samples was estimated for all genes and the top 100 up- and down-regulated genes ranked by differential promoter H3K4me3 were selected and tested for enriched GO terms and Reactome and PANTHER pathways using Enrichr (https://amp.pharm.mssm.edu). To determine if differences in H3K4me3 change were greater than expected by chance, average log likelihood for differential TBI versus sham H3K4me3 for DEGs for each gene expression trajectory module was compared against genes that were not DE via linear regression and ANOVA with Tukey’s post-hoc analysis were used to generate statistical significance results.

#### Data and Code Availability

The genomic data generated in this study and presented in this publication have been deposited in the NCBI database and are accessible through GEO Series accession number GSExxxxx (https://www.ncbi.nlm.nih.gov/geo/query/acc.cgi?acc=GSExxxxxx) and can be visualized in UCSC track hubs whose information is provided on Nord Lab GitHub page (https://nordneurogenomicslab.github.io/publications/).

## SUPPLEMENTARY MATERIAL

Supplementary Tables: Supplementary_Tables.xls

## REFERENCES

1. Centers for Disease Control and Prevention. (2019). Surveillance Report of Traumatic Brain Injury-related Emergency Department Visits, Hospitalizations, and Deaths—United States, 2014.

2. Defense and Veterans Brain Injury Center. (2019). DoD Numbers for Traumatic Brain Injury Worldwide — Totals 2019 Q1-Q4 (https://dvbic.dcoe.mil/sites/default/files/tbi-numbers/DVBIC_WorldwideTotal_2019.pdf). [cited 2020 Jun 25] Available from: https://dvbic.dcoe.mil/sites/default/files/tbi-numbers/DVBIC_WorldwideTotal_2019.pdf.

3. Masel, B.E., and DeWitt, D.S. (2010). Traumatic Brain Injury: A Disease Process, Not an Event. J. Neurotrauma 27, 1529–1540.

4. Crane, P.K., Gibbons, L.E., Dams-O’Connor, K., Trittschuh, E., Leverenz, J.B., Keene, C.D., Sonnen, J., Montine, T.J., Bennett, D.A., Leurgans, S., Schneider, J.A., and Larson, E.B. (2016). Association of Traumatic Brain Injury With Late-Life Neurodegenerative Conditions and Neuropathologic Findings. JAMA Neurol. 73, 1062–1069.

5. Chauhan, N.B. (2014). Chronic neurodegenerative consequences of traumatic brain injury. Restor. Neurol. Neurosci. 32, 337–365.

6. Gardner, R.C., and Yaffe, K. (2015). Epidemiology of mild traumatic brain injury and neurodegenerative disease. Mol. Cell. Neurosci. 66, 75–80.

7. Takahata, K., Tabuchi, H., and Mimura, M. (2016). Late-onset Neurodegenerative Diseases Following Traumatic Brain Injury: Chronic Traumatic Encephalopathy (CTE) and Alzheimer’s Disease Secondary to TBI (AD-TBI). Brain Nerve 68, 849—857.

8. Meng, Q., Zhuang, Y., Ying, Z., Agrawal, R., Yang, X., and Gomez-Pinilla, F. (2017). Traumatic Brain Injury Induces Genome-Wide Transcriptomic, Methylomic, and Network Perturbations in Brain and Blood Predicting Neurological Disorders. EBioMedicine 16, 184–194.

9. Cho, H., Hyeon, S.J., Shin, J.-Y., Alvarez, V.E., Stein, T.D., Lee, J., Kowall, N.W., McKee, A.C., Ryu, H., and Seo, J.-S. (2020). Alterations of transcriptome signatures in head trauma-related neurodegenerative disorders. Sci. Rep. 10, 8811.

10. Di Pietro, V., Amin, D., Pernagallo, S., Lazzarino, G., Tavazzi, B., Vagnozzi, R., Pringle, A., and Belli, A. (2010). Transcriptomics of traumatic brain injury: gene expression and molecular pathways of different grades of insult in a rat organotypic hippocampal culture model. J. Neurotrauma 27, 349–359.

11. Guo, X., Zhang, B., Gomez-Pinilla, F., Gao, F., and Zhao, Z. (2020). Secondary single-cell transcriptomic analysis reveals common molecular signatures of cerebrovascular injury between traumatic brain injury and aging. bioRxiv , 2020.06.29.178855.

12. Arneson, D., Zhang, G., Ying, Z., Zhuang, Y., Byun, H.R., Ahn, I.S., Gomez-Pinilla, F., and Yang, X. (2018). Single cell molecular alterations reveal target cells and pathways of concussive brain injury. Nat. Commun. 9, 3894.

13. Wong, V.S. (2016). Epigenetic changes following traumatic brain injury and their implications for outcome, recovery and therapy. Neurosci. Lett. 625, 26–33.

14. Nagalakshmi, B., Sagarkar, S., and Sakharkar, A.J. (2018). Epigenetic Mechanisms of Traumatic Brain Injuries. Prog. Mol. Biol. Transl. Sci. 157, 263–298.

15. Zhang, B., West, E.J., Van, K.C., Gurkoff, G.G., Zhou, J., Zhang, X.-M., Kozikowski, A.P., and Lyeth, B.G. (2008). HDAC inhibitor increases histone H3 acetylation and reduces microglia inflammatory response following traumatic brain injury in rats. Brain Res. 1226, 181–191.

16. McGuire, J.L., Ngwenya, L.B., and McCullumsmith, R.E. (2019). Neurotransmitter changes after traumatic brain injury: an update for new treatment strategies. Mol. Psychiatry 24, 995–1012.

17. Zhou, H., Chen, L., Gao, X., Luo, B., and Chen, J. (2012). Moderate traumatic brain injury triggers rapid necrotic death of immature neurons in the hippocampus. J. Neuropathol. Exp. Neurol. 71, 348–359.

18. Kawano, H., Kimura-Kuroda, J., Komuta, Y., Yoshioka, N., Li, H.P., Kawamura, K., Li, Y., and Raisman, G. (2012). Role of the lesion scar in the response to damage and repair of the central nervous system. Cell Tissue Res. 349, 169–180.

19. Felderhoff-Mueser, U., Schmidt, O.I., Oberholzer, A., Bührer, C., and Stahel, P.F. (2005). IL-18: a key player in neuroinflammation and neurodegeneration? Trends Neurosci. 28, 487–493.

20. Castranio, E.L., Mounier, A., Wolfe, C.M., Nam, K.N., Fitz, N.F., Letronne, F., Schug, J., Koldamova, R., and Lefterov, I. (2017). Gene co-expression networks identify Trem2 and Tyrobp as major hubs in human APOE expressing mice following traumatic brain injury. Neurobiol. Dis. 105, 1–14.

21. O’Brien, W.T., Pham, L., Symons, G.F., Monif, M., Shultz, S.R., and McDonald, S.J. (2020). The NLRP3 inflammasome in traumatic brain injury: potential as a biomarker and therapeutic target. J. Neuroinflammation 17, 104.

22. Luo, J., Yang, H., and Song, B.-L. (2020). Mechanisms and regulation of cholesterol homeostasis. Nat. Rev. Mol. Cell Biol. 21, 225–245.

23. Murray, S.C., Lorenz, P., Howe, F.S., Wouters, M., Brown, T., Xi, S., Fischl, H., Khushaim, W., Rayappu, J.R., Angel, A., and Mellor, J. (2019). H3K4me3 is neither instructive for, nor informed by, transcription. bioRxiv , 709014.

24. Ng, H.H., Robert, F., Young, R.A., and Struhl, K. (2003). Targeted Recruitment of Set1 Histone Methylase by Elongating Pol II Provides a Localized Mark and Memory of Recent Transcriptional Activity. Mol. Cell 11, 709–719.

25. Sato, M., Chang, E., Igarashi, T., and Noble, L.J. (2001). Neuronal injury and loss after traumatic brain injury: time course and regional variability. Brain Res. 917, 45–54.

26. Schmidt, R.H., Scholten, K.J., and Maughan, P.H. (1999). Time course for recovery of water maze performance and central cholinergic innervation after fluid percussion injury. J. Neurotrauma 16, 1139–1147.

27. Faden, A.I., Wu, J., Stoica, B.A., and Loane, D.J. (2016). Progressive inflammation-mediated neurodegeneration after traumatic brain or spinal cord injury. Br. J. Pharmacol. 173, 681–691.

28. Laitinen, T., Sierra, A., Bolkvadze, T., Pitkänen, A., and Gröhn, O. (2015). Diffusion tensor imaging detects chronic microstructural changes in white and gray matter after traumatic brain injury in rat. Front. Neurosci. 9, 128.

29. Bramlett, H.M., and Dietrich, W.D. (2002). Quantitative structural changes in white and gray matter 1 year following traumatic brain injury in rats. Acta Neuropathol. 103, 607–614.

30. Matzilevich, D.A., Rall, J.M., Moore, A.N., Grill, R.J., and Dash, P.K. (2002). High-density microarray analysis of hippocampal gene expression following experimental brain injury. J. Neurosci. Res. 67, 646–663.

31. Raghupathi, R. (2004). Cell death mechanisms following traumatic brain injury. Brain Pathol. 14, 215–222.

32. Laskowski, R., Creed, J., and Raghupathi, R. (2015). Pathophysiology of Mild TBI: Implications for Altered Signaling Pathways., in: Kobeissy, F. (ed). Brain Neurotrauma: Molecular, Neuropsychological, and Rehabilitation Aspects. Boca Raton (FL): CRC Press/Taylor & Francis.

33. Kokiko-Cochran, O., Ransohoff, L., Veenstra, M., Lee, S., Saber, M., Sikora, M., Teknipp, R., Xu, G., Bemiller, S., Wilson, G., Crish, S., Bhaskar, K., Lee, Y.-S., Ransohoff, R.M., and Lamb, B.T. (2015). Altered Neuroinflammation and Behavior after Traumatic Brain Injury in a Mouse Model of Alzheimer’s Disease. J. Neurotrauma 33, 625–640.

34. Karve, I.P., Taylor, J.M., and Crack, P.J. (2016). The contribution of astrocytes and microglia to traumatic brain injury. Br. J. Pharmacol. 173, 692–702.

35. Habgood, M.D., Bye, N., Dziegielewska, K.M., Ek, C.J., Lane, M.A., Potter, A., Morganti-Kossmann, C., and Saunders, N.R. (2007). Changes in blood-brain barrier permeability to large and small molecules following traumatic brain injury in mice. Eur. J. Neurosci. 25, 231–238.

36. Liu, Y., Beyer, A., and Aebersold, R. (2016). On the Dependency of Cellular Protein Levels on mRNA Abundance. Cell 165, 535–550.

37. Lipponen, A., Paananen, J., Puhakka, N., and Pitkänen, A. (2016). Analysis of Post-Traumatic Brain Injury Gene Expression Signature Reveals Tubulins, Nfe2l2, Nfkb, Cd44 and S100a4 as Treatment Targets. Sci. Rep. 6, 31570.

38. Wood, W.G., Li, L., Müller, W.E., and Eckert, G.P. (2014). Cholesterol as a causative factor in Alzheimer’s disease: a debatable hypothesis. J. Neurochem. 129, 559–572.

39. Barbero-Camps, E., Roca-Agujetas, V., Bartolessis, I., de Dios, C., Fernández-Checa, J.C., Marí, M., Morales, A., Hartmann, T., and Colell, A. (2018). Cholesterol impairs autophagy-mediated clearance of amyloid beta while promoting its secretion. Autophagy 14, 1129–1154.

40. Malik, B., Fernandes, C., Killick, R., Wroe, R., Usardi, A., Williamson, R., Kellie, S., Anderton, B.H., and Reynolds, C.H. (2012). Oligomeric amyloid-β peptide affects the expression of genes involved in steroid and lipid metabolism in primary neurons. Neurochem. Int. 61, 321–333.

41. Arenas, F., Garcia-Ruiz, C., and Fernandez-Checa, J.C. (2017). Intracellular Cholesterol Trafficking and Impact in Neurodegeneration. Front. Mol. Neurosci. 10, 382.

42. Cacabelos, R. (2017). Parkinson’s Disease: From Pathogenesis to Pharmacogenomics. Int. J. Mol. Sci. 18, 551.

43. Lemche, E. (2018). Early Life Stress and Epigenetics in Late-onset Alzheimer’s Dementia: A Systematic Review. Curr. Genomics 19, 522–602.

44. Wang, M., Wang, S., Li, Y., Cai, G., Cao, M., and Li, L. (2020). Integrated analysis and network pharmacology approaches to explore key genes of Xingnaojing for treatment of Alzheimer’s disease. Brain Behav. 10, e01610.

45. Sarnowski, C., Satizabal, C.L., DeCarli, C., Pitsillides, A.N., Cupples, L.A., Vasan, R.S., Wilson, J.G., Bis, J.C., Fornage, M., Beiser, A.S., DeStefano, A.L., Dupuis, J., and Seshadri, S. (2018). Whole genome sequence analyses of brain imaging measures in the Framingham Study. Neurology 90, e188 LP–e196.

46. Bezerra, G.A., Dobrovetsky, E., Seitova, A., Fedosyuk, S., Dhe-Paganon, S., and Gruber, K. (2015). Structure of human dipeptidyl peptidase 10 (DPPY): a modulator of neuronal Kv4 channels. Sci. Rep. 5, 8769.

47. Arneson, D., Zhang, Y., Yang, X., and Narayanan, M. (2018). Shared mechanisms among neurodegenerative diseases: from genetic factors to gene networks. J. Genet. 97, 795–806.

48. Pantaleo, N., Chadwick, W., Park, S.-S., Wang, L., Zhou, Y., Martin, B., and Maudsley, S. (2010). The mammalian tachykinin ligand-receptor system: an emerging target for central neurological disorders. CNS Neurol. Disord. Drug Targets 9, 627–635.

49. Ayka, A., and Şehirli, A.Ö. (2020). The Role of the SLC Transporters Protein in the Neurodegenerative Disorders. Clin. Psychopharmacol. Neurosci. 18, 174–187.

50. Chen, D., Zhu, J., Zhong, J., Chen, F., Lin, X., Dai, J., Chen, Y., Wang, S., Ding, X., Wang, H., Qiu, J., Wang, F., Wu, W., Liu, P., Tang, G., Qiu, X., Ruan, Y., Li, J., Zhu, S., Xu, X., Li, F., Liu, Z., and Cao, G. (2019). Single cell atlas of domestic pig brain illuminates the conservation and divergence of cell types at spatial and species levels. bioRxiv , 2019.12.11.872721.

51. Lalli, M.A., Garcia, G., Madrigal, L., Arcos-Burgos, M., Arcila, M.L., Kosik, K.S., and Lopera, F. (2012). Exploratory data from complete genomes of familial alzheimer disease age-at-onset outliers. Hum. Mutat. 33, 1630–1634.

52. Xicoy, H., Wieringa, B., and Martens, G.J.M. (2019). The Role of Lipids in Parkinson’s Disease. Cells 8, 27.

53. Van, K.C., and Lyeth, B.G. (2016). Lateral (Parasagittal) Fluid Percussion Model of Traumatic Brain Injury., in: Kobeissy, F.H., Dixon, C.E., Hayes, R.L., and Mondello, S. (eds). Injury Models of the Central Nervous System: Methods and Protocols. New York, NY: Springer New York, pps. 231–251.

54. Fedor, M., Berman, R.F., Muizelaar, J.P., and Lyeth, B.G. (2010). Hippocampal Theta Dysfunction after Lateral Fluid Percussion Injury. J. Neurotrauma 27, 1605–1615.

55. Gurkoff, G., Shahlaie, K., Lyeth, B., and Berman, R. (2013). Voltage-gated calcium channel antagonists and traumatic brain injury. Pharmaceuticals (Basel). 6, 788–812.

56. Lyeth, B.G., Jiang, J.I.Y.A.O., and Liu, S. (1993). Behavioral Protection by Moderate Hypothermia Initiated After Experimental Traumatic Brain Injury. J. Neurotrauma 10, 57–64.

57. Gompers, A.L., Su-Feher, L., Ellegood, J., Copping, N.A., Riyadh, M.A., Stradleigh, T.W., Pride, M.C., Schaffler, M.D., Wade, A.A., Catta-Preta, R., Zdilar, I., Louis, S., Kaushik, G., Mannion, B.J., Plajzer-Frick, I., Afzal, V., Visel, A., Pennacchio, L.A., Dickel, D.E., Lerch, J.P., Crawley, J.N., Zarbalis, K.S., Silverman, J.L., and Nord, A.S. (2017). Germline Chd8 haploinsufficiency alters brain development in mouse. Nat. Neurosci. 20.

58. Nord, A.S., Blow, M.J., Attanasio, C., Akiyama, J.A., Holt, A., Hosseini, R., Phouanenavong, S., Plajzer-Frick, I., Shoukry, M., Afzal, V., Rubenstein, J.L.R., Rubin, E.M., Pennacchio, L.A., and Visel, A. (2013). Rapid and Pervasive Changes in Genome-wide Enhancer Usage during Mammalian Development. Cell 155, 1521–1531.

59. Dobin, A., Davis, C.A., Schlesinger, F., Drenkow, J., Zaleski, C., Jha, S., Batut, P., Chaisson, M., and Gingeras, T.R. (2013). STAR: ultrafast universal RNA-seq aligner. Bioinformatics 29, 15–21.

60. Andrews, S. (2010). FASTQC. A quality control tool for high throughput sequence data. 2010. Http://Www.Bioinformatics.Babraham.Ac.Uk/Projects/Fastqc/ [cited 2019 Apr 16] Available from: http://www.bioinformatics.babraham.ac.uk/projects/fastqc.

61. Liao, Y., Smyth, G.K., and Shi, W. (2014). featureCounts: an efficient general purpose program for assigning sequence reads to genomic features. Bioinformatics 30, 923–930.

62. Ritchie, M.E., Phipson, B., Wu, D., Hu, Y., Law, C.W., Shi, W., and Smyth, G.K. (2015). limma powers differential expression analyses for RNA-sequencing and microarray studies. Nucleic Acids Res. 43, e47–e47.

63. Yu, G., Wang, L.-G., Han, Y., and He, Q.-Y. (2012). clusterProfiler: an R Package for Comparing Biological Themes Among Gene Clusters. Omi. A J. Integr. Biol. 16, 284–287.

64. Krueger, F. (2015). Trim Galore!: A wrapper tool around Cutadapt and FastQC to consistently apply quality and adapter trimming to FastQ files. Babraham Inst. Available from: http://www.bioinformatics.babraham.ac.uk/projects/trim_galore/.

65. Li, H., and Durbin, R. (2009). Fast and accurate short read alignment with Burrows–Wheeler transform. Bioinformatics 25, 1754–1760.

66. Li, H., Handsaker, B., Wysoker, A., Fennell, T., Ruan, J., Homer, N., Marth, G., Abecasis, G., Durbin, R., and Subgroup, 1000 Genome Project Data Processing. (2009). The Sequence Alignment/Map format and SAMtools. Bioinformatics 25, 2078–2079.

67. Zhang, Y., Liu, T., Meyer, C.A., Eeckhoute, J., Johnson, D.S., Bernstein, B.E., Nusbaum, C., Myers, R.M., Brown, M., Li, W., and Liu, X.S. (2008). Model-based Analysis of ChIP-Seq (MACS). Genome Biol 9, R137.

